# Efficacy of vocal cord injection of dedifferentiated fat cells in treating glottis closure insufficiency: Insights from a rat model of recurrent laryngeal nerve resection

**DOI:** 10.1101/2025.05.06.652489

**Authors:** Ikuo Mikoshiba, Kazuhiro Hagikura, Taro Matsumoto, Tomohiko Kazama, Ryohei Asai, Hiroyuki Hao, Takeshi Oshima

## Abstract

Glottic insufficiency results from impaired vocal cord contact, leading to a gap between the cords and manifesting as hoarseness and respiratory difficulties. Vocal cord injection is a commonly utilized therapeutic approach to rectify this gap by augmenting vocal cord volume; however, the optimal injectable material remains undetermined. Dedifferentiated fat cells (DFATs), derived from mature adipocytes, exhibit robust proliferative capacity and multipotency, establishing them as potential candidates for treating glottic insufficiency. This study investigated the therapeutic efficacy of DFATs in a rat model of recurrent laryngeal nerve paralysis. In the experimental design, the recurrent laryngeal nerve of each rat was resected unilaterally, and 5 weeks later, the atrophied vocal cord muscles were injected with either 10 μl of saline (control), 1.0 × 10^6^ DFATs in 10 μl (DFAT group), or 1.0 × 10^6^ DFATs combined with 500 ng of basic fibroblast growth factor (bFGF) in 10 μl (DFAT + bFGF group). At 4 weeks post-injection, laryngeal endoscopy was performed to evaluate the glottic gap, followed by histological analyses to assess vocal muscle atrophy, collagen deposition, and markers of cellular proliferation (Ki-67) and apoptosis (TUNEL). To confirm DFAT engraftment, GFP-labeled DFATs were evaluated at 2, 4, and 6 weeks post-injection. Results indicated that both the DFAT and DFAT + bFGF groups exhibited significantly reduced glottic gaps, increased collagen deposition, and decreased TUNEL-positive apoptotic cells compared to controls. Notably, the DFAT + bFGF group displayed superior outcomes, including a greater vocal muscle area and enhanced Ki-67-positive cell proliferation, indicating mitigation of muscle atrophy. GFP-positive DFATs were detectable in the tissue for up to 6 weeks, confirming engraftment. These findings underscore the potential of DFATs, particularly in combination with bFGF, as an innovative and effective therapeutic approach for glottic insufficiency secondary to recurrent laryngeal nerve paralysis.

## Introduction

Glottic insufficiency is a condition characterized by impaired contact between the vocal cords, resulting in the formation of a glottic gap that produces symptoms such as hoarseness and dyspnea. Pathologies contributing to glottic insufficiency include recurrent laryngeal nerve paralysis, vocal cord atrophy, vocal fold sulcus, and vocal cord scarring. In addition to surgical interventions such as thyroplasty and arytenoid adduction [1,2], intracordal injection therapy is employed for the treatment of glottic insufficiency. Intracordal injection refers to a technique in which a material is injected directly into the vocal cords via the oral cavity to augment their volume and correct their position inward, thereby reducing the glottic gap. This method is less invasive and more manageable than surgical procedures necessitating external cervical incisions; however, issues regarding the durability and stability of the effects have been reported [3]. Clinically utilized injectable materials for glottic insufficiency include autologous fat, atelocollagen, hyaluronic acid, and calcium phosphate paste [4–7]. Autologous fat is considered safer than other injectable materials, as it utilizes the patient’s own tissue and is highly effective for increasing vocal fold volume, leading to its widespread use since its introduction by Mikaelian et al. in 1991 [8].

In recent years, adipose-derived stem cells (ASCs) have garnered attention as a viable cell source for cell therapy. ASCs possess high proliferation and differentiation potential and can be harvested with minimal invasiveness from a small quantity of adipose tissue, compared to autologous fat. ASCs secrete humoral factors such as vascular endothelial growth factor (VEGF) and hepatocyte growth factor (HGF), which have been reported to promote wound healing, angiogenesis, and exhibit immunoregulatory effects [9]. The injection of ASCs into the vocal folds has been shown to inhibit scarring and enhance hyaluronic acid production in animal models of vocal fold injury, including studies in dogs, rats, and rabbits, resulting in improved healing [10–12]. Furthermore, combined injections of autologous fat and ASCs in cases of glottic insufficiency due to unilateral recurrent nerve paralysis in humans have been noted to improve fat graft survivorship, reduce the glottic gap, and enhance wound healing [13]. Thus, ASCs are anticipated to be effective materials for vocal fold injection procedures. However, the heterogeneity of ASCs, stemming from the incorporation of interstitial cell fractions extracted from tissue, is a noted limitation [14].

Dedifferentiated fat cells (DFATs) are cells characterized by high proliferation and multipotency, similar to ASCs, and they can be derived from mature adipocytes isolated via a method known as ceiling culture [15]. DFATs have been confirmed to possess the capability to differentiate into adipocytes, osteoblasts, chondrocytes, myofibroblasts, cardiac myocytes, skeletal myocytes, and pericytes [16–21]. Additionally, DFATs exhibit a secretion pattern of humoral factors similar to that of ASCs [22]. The isolation and culture of DFATs leverage the buoyant properties of mature adipocytes, providing the advantage of reduced contamination by foreign cells compared to ASCs and yielding high purity cells. Furthermore, a large number of cells can be generated from a small quantity (approximately 1 g) of aspirated fat, maintaining stable therapeutic activity regardless of the patient’s age or underlying conditions. Previous studies have demonstrated that the transplantation of DFATs in combination with basic fibroblast growth factor (bFGF) into a rat model of full-thickness skin defects resulted in improved vascular density and proliferation of dermis-like tissue [23]. bFGF has been clinically employed as an injectable material for treating glottic closure insufficiency, with therapeutic effects including voice improvement reported [24,25].

In this study, the potential for improving glottic closure insufficiency through the injection of DFAT into atrophied vocal cord muscles in a rat model of unilateral recurrent laryngeal nerve resection was explored. Additionally, the possibility of enhancing the therapeutic effect through concurrent administration of bFGF alongside DFATs was investigated.

## Materials and Methods

### Experimental Animals

Experiments were conducted using 8-week-old male Sprague-Dawley (SD) rats (Oriental Yeast Co., Ltd., Tokyo, Japan). This study adhered to the animal experiment protocol established by Nihon University School of Medicine. All animal procedures received approval from the Nihon University Animal Experiment Committee (approval numbers: AP20MED010, AP20MED011) and the Nihon University Genetic Recombination Experiment Safety Committee (approval number: 2019Medicine22). A total of 58 rats were used in this study. The experimental period lasted 7 to 11 weeks, depending on the specific protocol. Animals were monitored at least twice daily for clinical signs of pain, distress, or morbidity throughout the study. Humane endpoints were predefined, and euthanasia was performed when animals exhibited any of the following: significant weight loss (>20% of baseline), markedly reduced movement, diminished responsiveness, substantial reduction in food or water intake for more than 24 hours, or lack of grooming. Animals meeting these criteria were euthanized promptly via carbon dioxide (CO₂) inhalation, with death was confirmed by the absence of heartbeat and respiratory movement. During the study, two animals reached the criteria for humane euthanasia and were euthanized without delay. Three animals died during surgical procedures before meeting the predefined criteria; the cause of death was presumed to be anesthetic overdose. Analgesics and anesthetics were used as appropriate during invasive procedures to minimize pain and distress. Animals were housed in temperature- and humidity-controlled rooms under a 12-hour light/dark cycle, with ad libitum access to food and water. All animal care and handling were performed by trained personnel following approved institutional protocols.

### Cell culture

For this study, Green Fluorescent Protein (GFP)-labeled DFATs, derived from GFP transgenic rats [26], were utilized after being cryopreserved. The cryopreserved cells were thawed, cultivated in adherent culture, and subsequently used for experimentation. DFATs were maintained in tissue culture dishes (BD Falcon, Bedford, MA, USA) filled with Dulbecco’s modified Eagle’s medium (DMEM) (Thermo Fischer Scientific, Waltham, MA, USA) supplemented with 20% fetal bovine serum (FBS) (Sigma-Aldrich, St. Louis, MO, USA) and incubated at 37 ℃ in a 5% CO_2_ atmosphere. The culture medium was refreshed every 3 days until the cells reached confluence. GFP-positive cells were observed and imaged using a fluorescence microscope (BZ-X710, KEYENCE Corporation, Osaka, Japan).

### Creation of a rat model with unilateral recurrent laryngeal nerve paralysis

SD rats were anesthetized via intraperitoneal administration of medetomidine hydrochloride (0.15 mg/kg), midazolam (2 mg/kg), and butorphanol tartrate (2.5 mg/kg). Following hair removal from the incision site, a midline incision was made in the anterior neck while the rat was placed in the supine position. The anterior neck musculature was divided into left and right halves to expose both recurrent laryngeal nerves. Nerve manipulation was performed under a stereomicroscope (SZ2-ILST; OLYMPUS, Tokyo, Japan). After dissecting the connective tissue surrounding the trachea to verify the presence of both recurrent laryngeal nerves, the left recurrent laryngeal nerve was resected 10 mm at the level of the seventh tracheal ring, and both nerve ends were ligated with 5-0 Vicryl sutures. Post-procedure, the larynx of the rat was examined using a rigid endoscope (TrueView II; OLYMPUS), confirming the absence of left vocal cord movement and proper resection of the left recurrent laryngeal nerve.

### Intracordal injection

To establish a technique for intracordal injection in rats with unilateral recurrent laryngeal nerve paralysis, an injection needle and microsyringe for intracordal delivery were developed (Ito Works, Shizuoka, JAPAN). The injection needle featured two types of tip curvature, 30° and 45°, designed as a tapered needle that is 24 G in thickness halfway and 30 G only at the tip, minimizing interference with the endoscope during the injection process. After administering general anesthesia to SD rats, the submucosa of the anterior-lateral edge of the left arytenoid cartilage was punctured under visualization with a rigid endoscope (TrueView II; OLYMPUS), followed by the injection of experimental materials. Initially, 10 μl of collagen loaded with pyoktanin was injected employing the same technique. Immediately following the injection, the animals were euthanized using CO_2_ gas, and the larynx was excised for macroscopic examination. Macroscopic analysis confirmed that pyoktanin permeated from the lateral surface of the thyroid cartilage to the vocal cord muscle, indicating successful injection without leakage (S1 Fig.). Subsequently, 10 μl of saline solution containing the prepared GFP-labeled DFATs (1.0 × 10^6^ cells/10 μl) was injected into the vocal cords, and the animals were euthanized to prepare laryngeal tissue specimens. Within the laryngeal tissue, GFP-positive cell clusters were detected in the vocal cord muscle, confirming that DFATs had been accurately injected around the vocal cord muscle (S2 Fig.).

### Protocol 1 (Examination of therapeutic effects)

As previously described, 8-week-old male SD rats (n=25) underwent left recurrent laryngeal nerve resection, leading to the establishment of unilateral recurrent laryngeal nerve paralysis model. The rats were thereafter allocated into three groups (n = 8–9 rats per group): (i) control group (saline), (ii) DFAT group, and (iii) DFAT + bFGF group. Five weeks following nerve resection, the control group received 10 μl of saline, the DFAT group received DFAT (1.0×10^6^ cells / 10 μl of saline), and the DFAT + bFGF group received DFAT (1.0×10^6^ cells) combined with bFGF (500 ng /10 μl of saline). bFGF was utilized by dissolving Fiblast^TM^ Spray 250 (Kaken Pharmaceutical Co.), which contains 250 μg of lyophilized human recombinant bFGF, in 5 mL of saline immediately before administration. Rats under general anesthesia were positioned on an intubation table (KN-1014; Natsume Seisakusho Co., Ltd., Tokyo, Japan) for injection into the left vocal cord, using a rigid endoscope for observation.

While under general anesthesia, rats were secured on an intubation table, facilitating the observation of laryngeal movement through a rigid endoscope while the tongue was drawn forward. Evaluations occurred immediately prior to the injection procedure and 4 weeks thereafter. The rat larynx is characterized by small vocal cords and large arytenoid cartilages, necessitating an evaluation methodology that emphasizes the mobility of the arytenoid cartilage. The adduction angle of the arytenoid cartilage was analyzed using image analysis software (ImageJ, ver. 1.51; National Institutes of Health, Rockville, MD), identifying the angle formed by the midline of the glottis and a tangent drawn from the junction of the arytenoid cartilages on both sides to the left (treated) arytenoid cartilage. The adduction angle of the arytenoid cartilage and the improvement rate of the glottic gap were measured just before and 4 weeks after injection as follows (S3 Fig.).

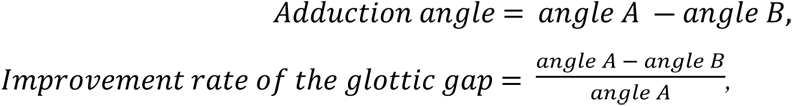

where, angle A represents the degree of separation between the vocal folds pre-treatment; and angle B represents the angle of the glottic gap post-treatment.

At 4 weeks post-treatment, all rats were euthanized using CO_2_ gas, and the larynges were excised for specimen preparation. Histological examination of the vocal cord muscle involved assessing its cross-sectional area and the extent of collagen accumulation therein. The excised larynges were fixed in 10% neutral buffered formalin (Fujifilm, Tokyo, Japan), embedded in paraffin, and sectioned at 5-μm thickness at the level where the arytenoid cartilage was visible. Tissue sections were stained with hematoxylin and eosin (HE) and Masson’s trichrome stain, then observed and photographed using a fluorescence microscope (BZ-x710; KEYENCE). Sections exhibiting the most symmetrical shapes of the left and right arytenoid cartilages were selected for evaluation. The cross-sectional area of the vocal cord muscle was determined by tracing the circumference of the vocal cord muscle using ImageJ and converting it into two colors on optical micrographs of HE stained samples. The cross-sectional area of the muscle layer on the treated side was measured relative to the normal (untreated) side, and the cross-sectional area ratio (T/U ratio) was calculated. To evaluate collagen accumulation, the outer circumference of the vocal cord muscle was traced using ImageJ on optical micrographs of Masson’s trichrome-stained samples; the area of blue-staining collagen within the muscle layer was extracted, and the area ratio (T/U) was computed.

### Protocol 2 (Study of mechanism of action)

A recurrent laryngeal nerve paralysis model was generated in 8-week-old male SD rats (n=25), assigned to three groups (n=8–9 each). Two weeks after injection, all rats were euthanized, and the larynges were removed for specimen preparation. Apoptotic cells were evaluated through TUNEL-positive cell detection, while proliferative cells were assessed via Ki-67-positive cell identification. For immunohistochemical examination of the vocal cord muscle, the excised larynges were fixed in 10% neutral buffered formalin, embedded in paraffin, and cut into 5-μm thick sections at the level where the arytenoid cartilage was discernible. Following deparaffinization, samples underwent antigen retrieval by immersion in target retrieval solution (DakoCytomation, Glostrup, Denmark) and were autoclaved for 15 minutes. After rewarming, sections were treated with blocking buffer (PBS containing 1% BSA and 0.5% Triton X-100) for 1 hour to inhibit nonspecific reactions. After washing with PBS, the sections were incubated overnight at 4 °C with rabbit anti-Ki-67 antibody (1:100; Abcam, Cambridge, MA, USA) as the primary antibody. The following day, sections were washed twice with PBS and incubated at room temperature for 1 hour with Alexa Fluor 488 Donkey anti-rabbit IgG (H+L) antibody (1:400; Invitrogen, Carlsbad, CA, USA) as the secondary antibody. Following PBS washes, sections were incubated at room temperature for 15 minutes with Hoechst 33342 (1:1000; Invitrogen). After washing with distilled water, sections were mounted with ProLong™ Gold antifade mountant (Invitrogen). The DeadEnd™ Fluorometric TUNEL system (Promega, Madison, WI, USA) was utilized to detect TUNEL-positive cells. All stained specimens were analyzed and photographed using a fluorescence microscope or a confocal laser scanning microscope (FLUOVIEW FV10i; OLYMPUS). For TUNEL-positive and Ki-67-positive cells, a composite image of the entire cross-section of the vocal cord muscle was captured, and the entire circumference of the vocal cord muscle was traced to calculate the quantity of each positive cell type present within the vocal cord muscle.

### Protocol 3 (Examination of DFATs engraftment)

A recurrent laryngeal nerve paralysis model was established in 8-week-old male SD rats (n=3) utilizing the aforementioned procedures, with all rats receiving DFATs (1.0×10^6^ cells/10 μl of saline) 5 weeks following nerve resection. Post-injection, the rats were euthanized at 2, 4, and 6 weeks (n=1 each), and the larynges were excised for specimen preparation. GFP-positive cells were evaluated to confirm DFAT engraftment. Post-washing with PBS, sections were incubated with Goat anti-GFP antibody (1:100; Abcam) as the primary antibody and Alexa Fluor 647 Donkey anti-goat IgG (H+L) antibody (1:400; Invitrogen) as the secondary antibody. Following PBS washes, sections were incubated with Hoechst 33342 (1:1000; Invitrogen) for 15 minutes at room temperature. The sections were thoroughly washed with distilled water and mounted with ProLong™ Gold antifade mountant (Invitrogen). Observations of GFP-positive cells were conducted using a fluorescence microscope or a confocal laser scanning microscope (FLUOVIEW FV10i; OLYMPUS).

### Statistical analysis

Quantitative data obtained from the experiments were statistically analyzed using a one-way analysis of variance (one-way ANOVA), with Tukey’s multiple comparisons test employed as a post-hoc procedure. A p-value of <0.05 was deemed statistically significant (GraphPad Prism ver 8.0, La Jolla, CA, USA).

## Results

To assess the efficacy of interventions for improving glottic closure insufficiency across groups, the adduction angle of the arytenoid cartilage and the glottic gap improvement rate were measured using a laryngeal endoscope. The change in the adduction angle (mean ± standard deviation) from baseline to 4 weeks post-injection was significantly greater in both the DFAT group (2.7 ± 1.5°) and the DFAT + bFGF group (2.5 ± 1.3°) compared to the control group (0.06 ± 0.45°) (Fig. 1; p < 0.001 for both). In addition, the glottic gap improvement rate was significantly higher in the DFAT group (25.1 ± 9.1%) and the DFAT + bFGF group (28.2 ± 13.8%) relative to the control group (0.8 ± 6.3%; p < 0.001 for both). No significant difference was observed in the improvement rates for the adduction angle and the glottic gap between the DFAT and DFAT + bFGF groups.

**Fig 1.**
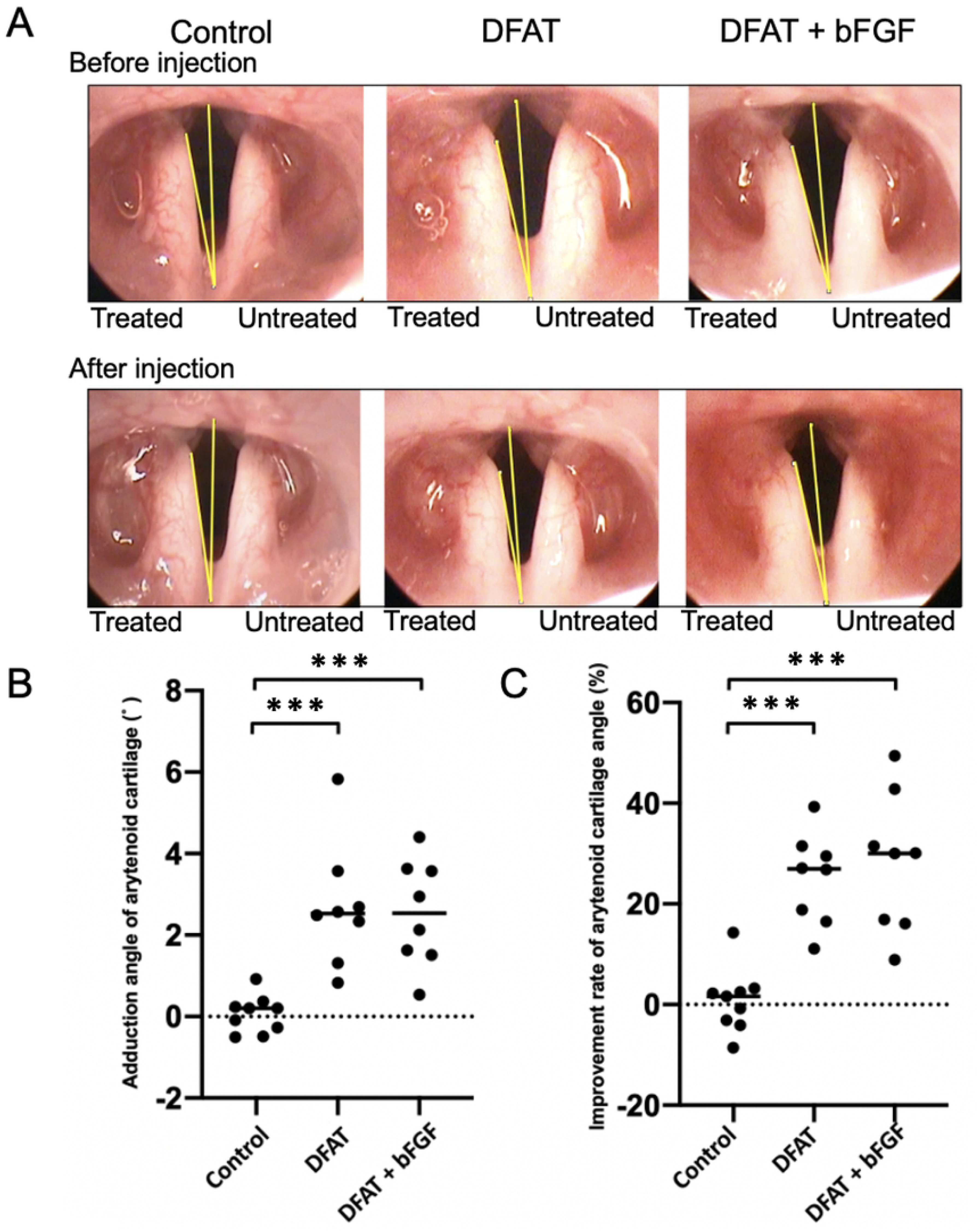
Effect of DFATs transplantation in improving the glottis gap. At 5 weeks following unilateral recurrent nerve transection, injections of saline, DFATs, and DFATs combined with bFGF (DFAT + bFGF) were administered into the vocal folds. At 4 weeks post-injection, the adduction deviation angle of the arytenoid cartilage and the improvement rate of the glottic gap in each group were evaluated via laryngoscopy. **(A)** Representative laryngoscopy images of each group. **(B)** Change in adduction deviation angle from baseline to 4 weeks post-injection. **(C)** Improvement rate of arytenoid cartilage angle from baseline to 4 weeks post-injection. These findings indicate that the injection of DFATs and DFATs + bFGF into atrophied vocal fold muscles effectively minimized the glottic gap. ***p < 0.001 (one-way ANOVA, Tukey’s multiple comparisons test).

To evaluate the effect on vocal cord muscle atrophy in each group, the extent of atrophy of the vocal cord muscle was assessed using HE-stained images obtained 4 weeks after intracordal injection. The T/U ratio of the vocal cord muscle cross-sectional area (mean ± standard deviation) was significantly higher in the DFAT + bFGF group (0.91 ± 0.18) than in the control group (0.69 ± 0.14) (Fig. 2; p < 0.05 for both). The T/U ratio in the DFAT group (0.81 ± 0.11) demonstrated a trend toward being higher than that of the control group; however, this difference was not statistically significant. These findings indicate that the combined injection of DFATs and bFGF into atrophied vocal fold muscles effectively inhibits muscle atrophy.

**Fig 2.**
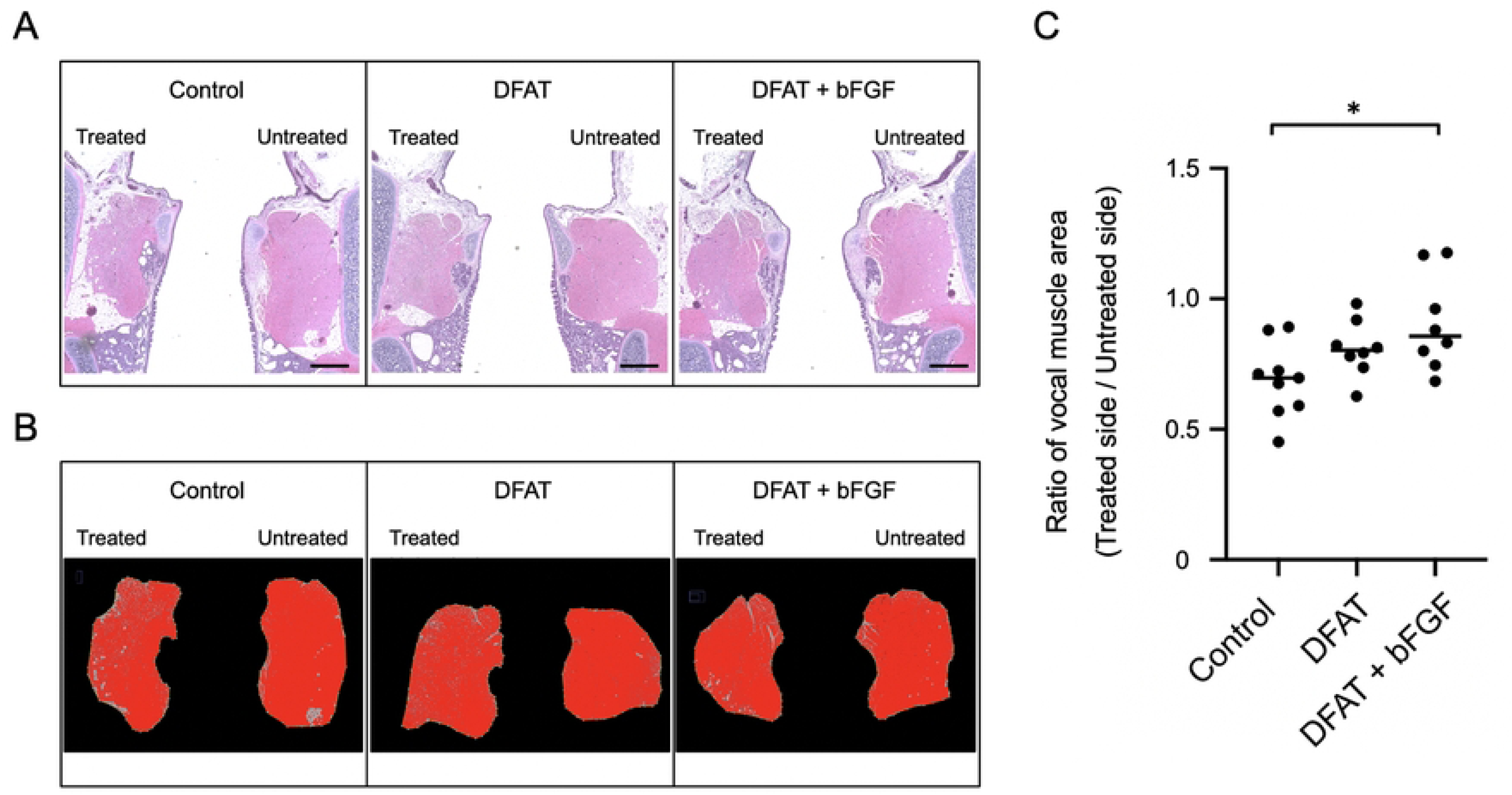
Effect of DFATs implantation on the inhibition of atrophy in the atrophic vocal fold muscle. At 5 weeks following unilateral recurrent nerve transection, injections of saline, DFATs, and DFATs + bFGF were administered into the vocal folds, and the extent of vocal fold muscle atrophy in each group was evaluated 4 weeks after injection using H&E staining. **(A)** Representative image of the vocal fold muscle (H&E staining) following injection in each group. Scale bar: 500 μm. **(B)** Quantification of the cross-sectional area of the vocal fold muscle (red) using ImageJ. **(C)** T/U ratio of the cross-sectional area of the vocal fold muscle in each group. * p < 0.05 (one-way analysis of variance, Tukey’s multiple comparisons test).

To compare collagen accumulation in the vocal fold muscles across groups, the collagen content in the treated vocal fold muscles was assessed 4 weeks after intracordal injection using Masson’s Trichrome staining. The T/U ratio (mean ± standard deviation) of collagen accumulation in the vocal fold muscles was significantly greater in the DFAT group (3.72 ± 1.70; p < 0.01) and the DFAT + bFGF group (3.43 ± 0.96; p < 0.05) compared to the control group (1.66 ± 0.82) (Fig. 3). No significant difference was detected between the DFAT and DFAT + bFGF groups. These findings suggest that both DFAT injection and the combined injection of DFATs and bFGF into atrophic vocal fold muscles effectively promoted collagen accumulation in the vocal fold muscles.

**Fig 3.**
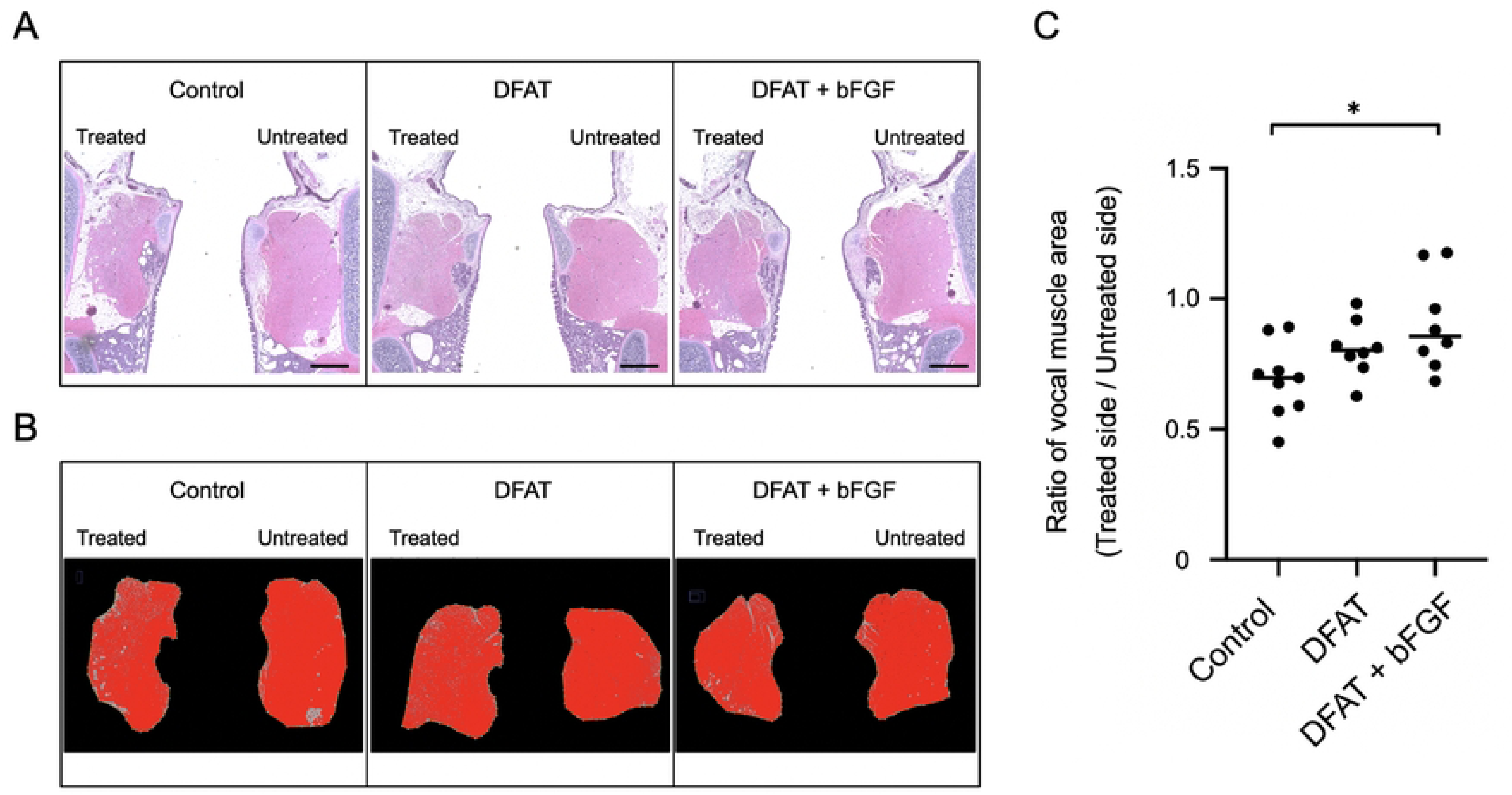
Effect of DFATs transplantation on collagen accumulation within the atrophic vocal fold muscle. At 5 weeks following unilateral recurrent nerve transection, injections of saline, DFATs, and DFATs + bFGF were administered into the vocal folds, and collagen accumulation within the vocal fold muscle of each group was evaluated 4 weeks post-injection using Masson’s Trichrome staining. **(A)** Representative image of the vocal fold muscle (Masson’s Trichrome staining) after injection for each group. Scale bar: 500 μm. **(B)** Quantification of collagen accumulation within the muscle layer using Image J. **(C)** T/U ratio of collagen accumulation within the vocal fold muscle in each group. * p < 0.05, ** p < 0.01 (one-way analysis of variance, Tukey’s multiple comparisons test).

To compare cell proliferation in atrophic vocal fold muscle tissue across groups, Ki-67 positive cells were quantified in the vocal fold muscle tissue 2 weeks after treatment. On microscopic examination, Ki-67 positive cells were primarily observed within the interstitial regions (Fig. 4A). The number of Ki-67 positive cells (mean ± SD) in the treated vocal fold muscle was significantly higher in the DFAT + bFGF group (25.0 ± 9.5) compared to the control group (9.4 ± 6.2; p < 0.01) and the DFAT group (13.5 ± 8.9; p < 0.05). No significant difference was observed between the control group and the DFAT group (Fig. 4B). These findings indicate that the combined injection of DFATs and bFGF into the atrophied vocal fold muscle promotes the induction of Ki-67 positive cells.

**Fig 4.**
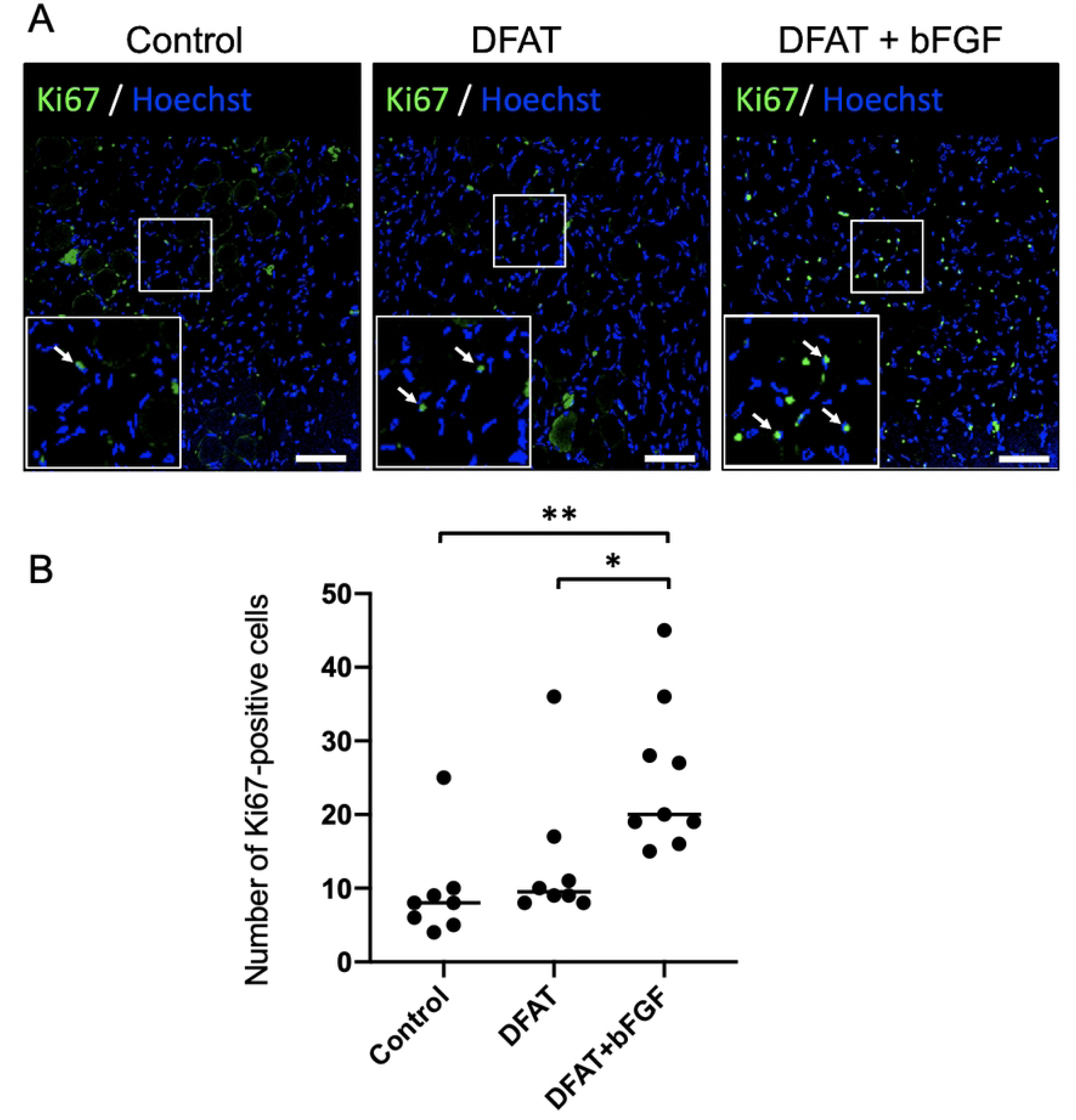
Assessment of cell proliferation within the vocal fold muscle following DFATs transplantation. At 5 weeks following unilateral recurrent nerve transection, injections of saline, DFATs, and DFATs + bFGF were administered into the vocal folds, and Ki-67 positive cells within the vocal fold muscle of each group were assessed 2 weeks after injection. **(A)** Representative fluorescent immunohistology images for each group (arrows indicate Ki-67 positive cells). Scale bars: 400 μm. **(B)** Number of Ki-67 positive cells in each group. * p < 0.05, ** p < 0.01 (one-way analysis of variance, Tukey’s multiple comparisons test).

To assess apoptosis in the atrophied vocal fold muscle tissue across groups, TUNEL-positive cells were quantified in the vocal fold muscle tissue at 2 weeks following treatment. The number of TUNEL-positive cells (mean ± standard deviation) in the treated vocal fold muscle was significantly lower in both the DFAT group (17.9 ± 8.1; p < 0.05) and the DFAT + bFGF group (18.0 ± 3.5; p < 0.01) compared to the control group (31.8 ± 8.3) (Fig. 5). No significant difference was detected between the DFAT and DFAT + bFGF groups. These findings demonstrate that the injection of DFATs, as well as the combined injection of DFATs and bFGF into the atrophied vocal fold muscle, effectively inhibits apoptosis in these tissues.

**Fig 5.**
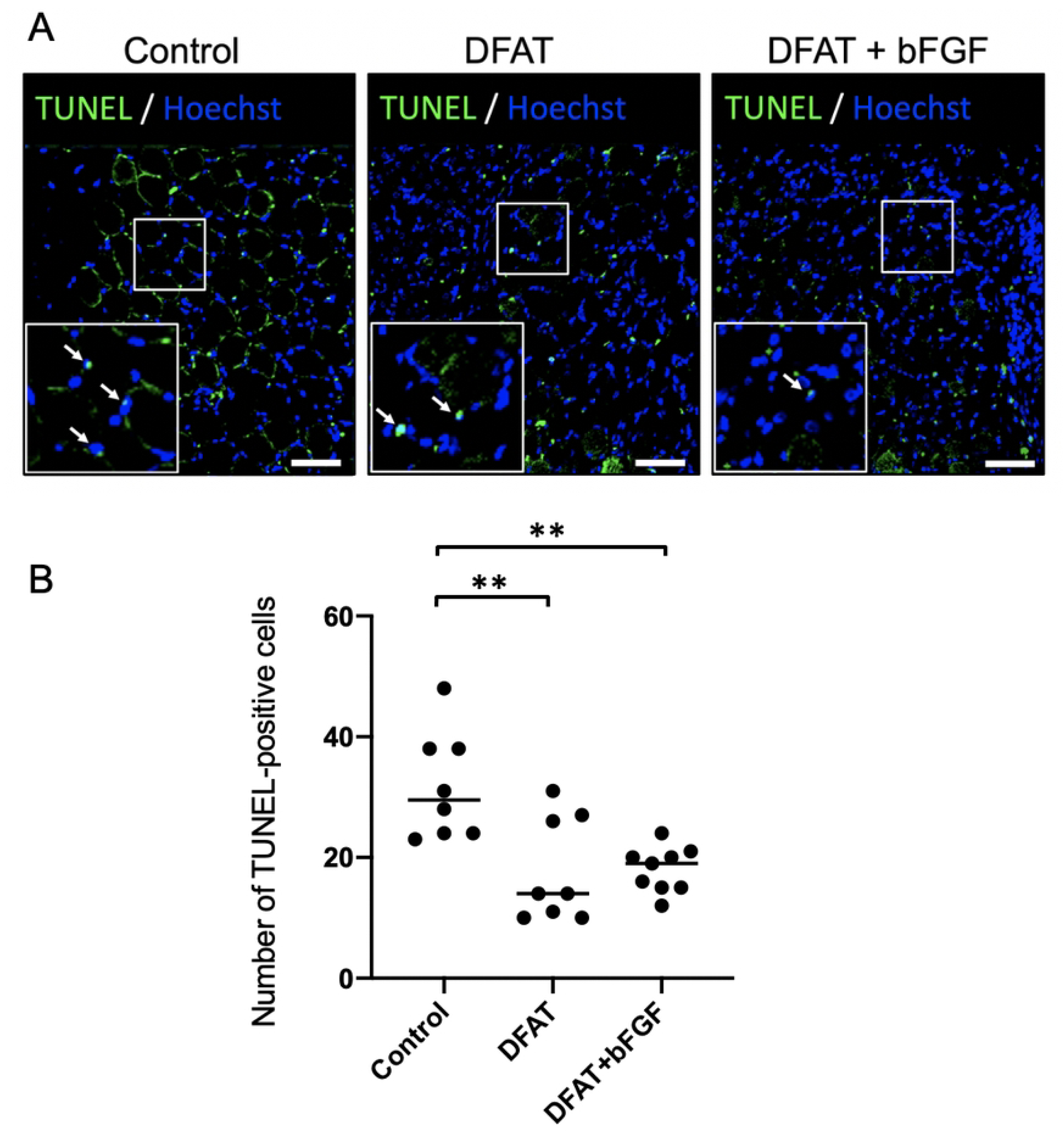
Analysis of apoptotic cells within the vocal fold muscle following DFATs transplantation. At 5 weeks following unilateral recurrent nerve transection, saline, DFATs, and DFATs + bFGF were injected into the vocal folds, and TUNEL-positive cells within the vocal fold muscle of each group were assessed 2 weeks post-injection. **(A)** Representative fluorescent immunohistological images for each group (arrows indicate TUNEL-positive cells). Scale bars: 400 μm. **(B)** Number of TUNEL-positive cells within the vocal fold muscle on the treatment side. ** p<0.01 (one-way analysis of variance, Tukey’s multiple comparisons test).

To assess the survival of transplanted DFATs in the vocal fold muscle, histological evaluation of the vocal fold tissue was performed at 2, 4, and 6 weeks following the transplantation of GFP-positive DFATs. At 2 weeks post-transplantation, numerous GFP-positive cells were observed within the interstitium of the transplanted vocal fold muscle (Fig. 6). While GFP-positive cells remained detectable at both 4 and 6 weeks post-transplantation, their numbers diminished significantly. These findings confirm that DFATs transplanted into atrophic vocal fold muscle survive for at least 6 weeks.

**Fig 6.**
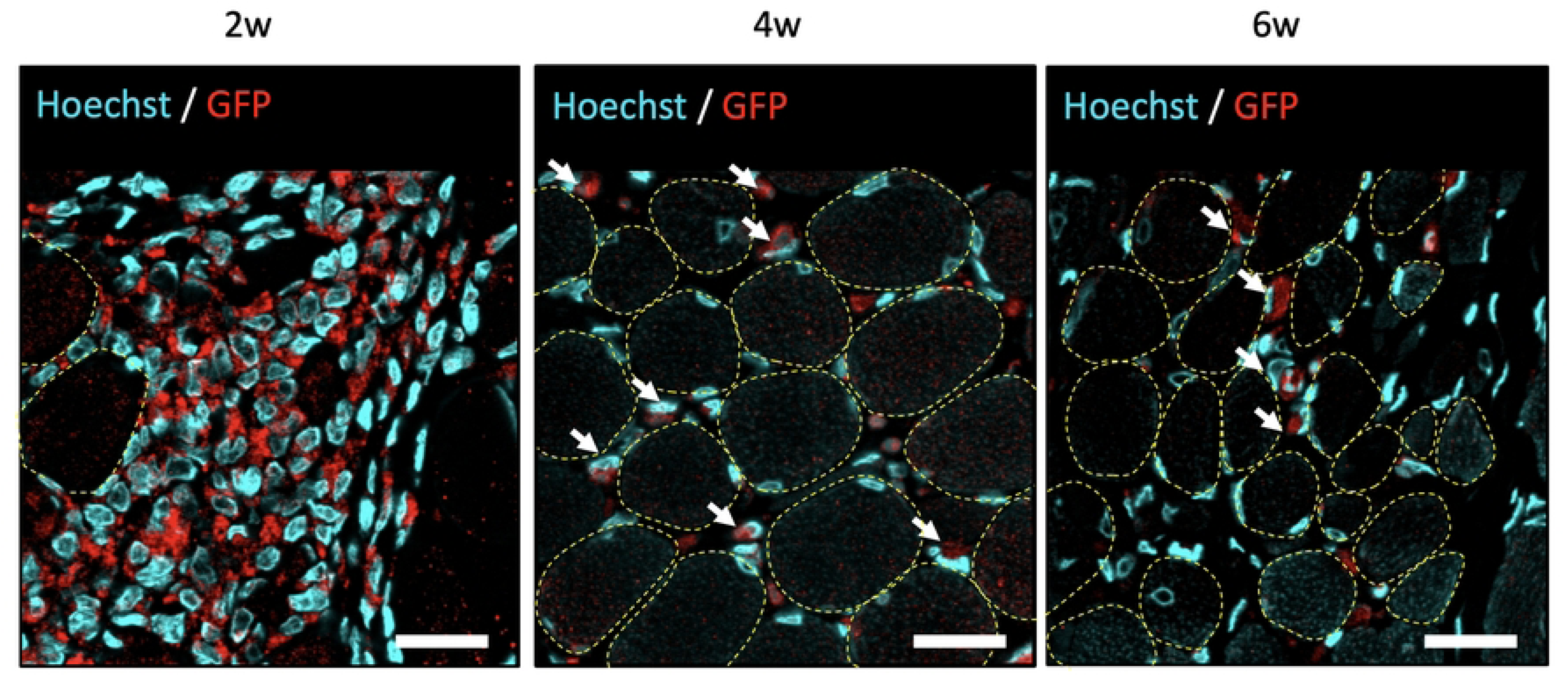
Assessment of the viability of DFATs within the vocal fold muscle using GFP immunostaining. At 5 weeks following unilateral recurrent nerve transection, GFP-labeled DFATs were injected into the vocal folds, and GFP-positive cells within the vocal fold muscle of each group were assessed at 2, 4, and 6 weeks post-injection. Fluorescence immunohistology images of the vocal fold muscle are presented (arrows indicate GFP-positive cells; dotted line indicates myocytes). Scale bars: 20 μm.

## Discussion

In this study, DFATs or combination of DFATs and bFGF were administered to the atrophied vocal cord muscles of rats following the resection of the recurrent laryngeal nerve. The therapeutic effects on vocal cord muscle regeneration and the reduction of the glottic gap were systematically evaluated. Laryngeal endoscopic assessments revealed a significant decrease in the glottic gap, which was attributed to the transplantation of DFATs into the atrophied vocal cord muscles. This work represents the first report highlighting the potential utility of cell therapy in addressing glottic gap resulting from recurrent laryngeal nerve paralysis.

Histological examination indicated that DFAT transplantation significantly reduced the presence of apoptotic cells in the vocal cord muscles while promoting collagen accumulation. It is believed that these effects contributed to the observed reduction in the glottic gap. Moreover, a similar decrease in apoptotic cells following DFAT transplantation has been documented in a rat model of stress urinary incontinence, whereby DFATs were introduced into the urethral sphincter [27]. Previous studies have highlighted that DFATs secrete humoral factors, such as VEGF and HGF, which play critical roles in cell survival and angiogenesis [21]. Based on these findings, it is plausible that the paracrine activity of the humoral factors secreted by the transplanted DFATs contributed to the reduction in apoptotic cells observed in the vocal fold muscles in this study.

Masson’s Trichrome staining revealed a significant increase in collagen accumulation between the muscle bundles of the atrophied vocal cord muscles following DFAT transplantation. Previous research has shown that DFATs secrete transforming growth factor-beta (TGF-β), a key regulator of collagen synthesis [21]. Additionally, it has been reported that DFATs can differentiate into myofibroblasts, which produce substantial amounts of collagen when stimulated by TGF-β [19]. Furthermore, the transplantation of DFATs into full-thickness skin wounds in pigs has been shown to enhance collagen deposition in the dermis [23]. The observed increase in collagen accumulation after DFAT transplantation may be attributed to two potential mechanisms: the paracrine action of TGF-β secreted by DFATs, which stimulates collagen production in surrounding tissues, or the transformation of transplanted DFATs into myofibroblasts, thus directly enhancing collagen secretion. Furthermore, the co-administration of bFGF alongside DFAT transplantation effectively inhibited vocal fold muscle atrophy and resulted in a significant increase in Ki-67-positive cells, indicating enhanced cellular proliferation.

bFGF is known to be involved in the proliferation, differentiation, and migration of various cell types. A previous study reported that the injection of bFGF into atrophic vocal fold muscle in a rat model of recurrent laryngeal nerve paralysis promoted myoblast proliferation while suppressing muscle atrophy [28]. Based on these findings, it can be inferred that the concomitant administration of bFGF in this study stimulated cellular proliferation within the atrophic vocal fold muscles, thereby mitigating muscle atrophy.

In this study, Ki-67-positive cells were predominantly observed in the perimuscular regions and the interstitial spaces, with their morphology suggesting they were muscle satellite cells or fibroblasts. Another potential mechanism underlying the suppression of vocal fold muscle atrophy due to the combined administration of DFATs and bFGF may involve the synergistic effects of bFGF and humoral factors secreted by DFATs, such as VEGF and HGF, which may enhance angiogenesis and tissue repair. It is further hypothesized that the administered bFGF stimulated DFATs, thereby enhancing their proliferative capacity and therapeutic efficacy. However, additional investigations are warranted to elucidate the precise mechanisms by which the combined administration of DFATs and bFGF amplifies therapeutic outcomes.

Assessment of GFP-labeled DFAT engraftment confirmed that transplanted DFATs remained viable at the transplantation site for at least 6 weeks. The GFP-positive cells were primarily localized within the interstitium, with their numbers significantly declining by the fourth week. This suggests that the transplanted DFATs were unlikely to directly differentiate into vocal cord muscle tissue or contribute to the improvement of the glottic gap through a bulking effect.

This study had certain limitations. First, improvements in the glottic gap were observed up to 4 weeks following DFAT transplantation; this limited timeframe raises questions about the sustainability of the observed improvements in glottic closure insufficiency and whether the benefits persist beyond the short-term. Further investigation is essential to determine whether this therapeutic effect is maintained over a longer observation period. Second, the engraftment studies assessing the persistence of GFP-positive DFATs were conducted on only three rats, which may not provide a statistically robust representation of the efficacy and survival of transplanted DFATs across a wider population. Increasing the sample size in future engraftment studies would enhance statistical power, ensure more reliable data, and provide a better understanding of the behavior of DFATs over time. Additionally, future studies should explore optimal transplantation conditions for DFATs to achieve prolonged and consistent therapeutic benefits.

## Conclusion

In summary, the injection of DFATs into the atrophied vocal cord muscles of a rat model of unilateral recurrent laryngeal nerve resection effectively improved glottic closure insufficiency. Furthermore, the co-administration of bFGF demonstrated an ability to mitigate vocal cord muscle atrophy and stimulate cellular proliferation. The administration of DFATs, either alone or in combination with bFGF, holds promise as a novel therapeutic approach for addressing glottic closure insufficiency.

## Acknowledgements

We would to thank Editage (www.editage.jp) for English language editing.

## Supporting Information

**S1 Fig. Endoscopic injection into the vocal fold muscle.**

After general anaesthesia was administered to Sprague-Dawley (SD) rats, 10 µl of collagen gel containing pioctanine dye was injected into the vocal fold muscle, using a custom-made injection needle and micro syringe, under the guidance of a rigid endoscope (TrueView II; OLYMPUS). After injection, the rats were euthanized, and the larynges were excised for examination.

**(A)** Endoscopic view of the intralaryngeal vocal fold muscle injection procedure. **(B)** Rigid endoscopic image during vocal fold injection. Dotted line indicates the injection needle; arrow points to the injection site. **(C)** Rigid endoscopic image immediately after injection. **(D)** Appearance of the larynx after laryngectomy.

**S2 Fig. Injection and evaluation of GFP-labelled DFATs in vocal fold muscle.**

GFP-labelled DFATs mixture (10 µl) was injected endoscopically into the vocal fold muscles of the rats; and immediately afterwards the rats were euthanized and the larynges were excised. Frozen specimens were then prepared for immunohistological evaluation.

**(A)** DFATs during culture. Scale bar = 300 µm. **(B)** Fluorescence immunostained image of laryngeal tissue post-transplantation. Scale bar = 200 μm. Arrow: injection point; dotted line: muscle area.

**S3 Fig. Methods of assessment of the glottic gap using a rigid endoscope**

Vocal fold movement was video-recorded under laryngeal endoscopy, and the angle formed by the midline of the glottis and a tangent drawn from the junction of the arytenoid cartilages on both sides to the left (treated) arytenoid cartilage was measured before injection (angle A) and 4 weeks after injection (angle B), respectively. The adduction angle (angle A - angle B) and the improvement rate of the glottic gap ([angle A - angle B]/angle A) were then calculated.

